# Social communication between microbes colonizing the social honey bee *Apis mellifera*

**DOI:** 10.1101/287995

**Authors:** K.I. Miller, C.D. Franklin, H. R. Mattila, I.L.G. Newton

## Abstract

The European honey bee (*Apis mellifera*) is a charismatic species that plays a critical role in the pollination of agriculturally important crops and native flora. One emerging field of research is that of the host-associated honey bee microbiome: a group of bacterial phylotypes consistently found within the honey bee, which may play critical roles such as protection from pathogens and nutrient acquisition. In other model systems, host-associated microbial communities are known to participate in a form of bacterial communication known as quorum sensing. This type of communication allows bacteria to sense their environment and respond with changes in gene expression, controlling a number of factors including virulence, biofilm formation, and cell motility. Here, we have investigated the production of a specific quorum sensing molecule by honey bee microbes *in vivo* and *in vitro*. We specifically focused on the inter-species signaling molecule, autoinducer-2 (AI-2). We identified the production of AI-2 by both the entire community (using honey bee gut homogenates) and by cultured isolates, using a *Vibrio harveyi* biosensor. By comparing newly emerged and adult bees, we showed this signal is likely coming from the core microbial community. Finally, using honey bee specific bacterial isolates, we identified changes in biofilm production when isolates are exposed to increased levels of exogenous AI-2. Altogether, these data provide multiple lines of evidence for the presence of quorum sensing inside the honey bee host. The effect of AI-2 on biofilm formation by honey bee specific bacteria identifies one potential avenue for quorum sensing to affect host health.

**Author summary:** Microbial communities associate with every animal on the planet and can have dramatic effects on the health of their host. The honey bee is one such animal, home to a characteristic community of bacteria, which may provide various benefits. Here, we show that these microbes are producing quorum sensing molecules which could support interactions between bacterial members and facilitate host colonization.

## Introduction

Host-associated microbial communities (the microbiota) can have dramatic effects on the health, fecundity, and longevity of many insect hosts. For example, germ free *Drosophila* are unable to survive the larval stage when in low nutrient environments and their survival can be restored with the addition of just one bacterial strain (1, 2). Additionally, alterations in the microbiome can affect traits as from mating behavior (3) to protection from pathogens (4, 5). These examples point to the critical and varied roles of the microbiota in insects. Insects are the most numerically and taxonomically abundant animal group on the planet and play important roles in disease ecology (6), herbivory (7), pollination (8), and other ecosystem processes (9, 10). It is therefore vital that we understand how insect associated microbes may shape insect health and subsequently or directly, impact their ecological roles.

The honey bee gut microflora is described as a consistent group of bacterial clades, dominated by Gamma-proteobacteria, Firmicutes, and Actinobacteria (11–14). Basic characterization of these microbial groups has led to speculations about their role in honey bee health and whether they are responsible for provisioning nutrients (15) or assisting in the breakdown of plant-derived carbohydrates (16, 17), as is the case for other insect-associated microbes. Additionally, honey bees are more susceptible to pathogens after their microbiome is disrupted by antibiotics, supporting a protective role of the microbiota (5). Although we are just starting to understand the functions of these microbial species, we do know that they interact with each other *in vivo* (18) and *in vitro* (19). For example, *Gilliamella, Snodgrassella*, and *Lactobacillus* strains together form a biofilm on host tissue in the ileum of honey bees (18). When grown in co-culture, lactic acid bacteria found in the honey bee promote each other’s growth, suggesting a mutualistic or syntrophic interaction (19). However, little is currently known about *how* the honey bee associated microbes interact with each other and how these interactions impact the host.

It is important to emphasize that bacterial species do not exist in isolation; although studied in monoculture in the laboratory, their natural ecology includes other microbial organisms. One way bacteria communicate in their natural ecology is through a process referred to as “quorum sensing”. Quorum sensing is broadly phylogenetically conserved, found in a variety of bacterial classes (20). This density-dependent communication allows bacteria to sense their environment and make population scale behavioral changes in response. Bacteria participating in quorum sensing produce signaling molecules, termed autoinducers, and the concentration of these molecules correlates with bacterial density. When a threshold is reached, autoinducers often elicit changes in gene expression which affect many processes including biofilm formation, symbiosis, motility, virulence and a number of others (21, 22). For example, the bobtail squid forms an intimate, mutualistic relationship with *Vibrio fisheri*, which produces bioluminescence at high densities, providing beneficial camouflage for the host (28). In contrast, quorum sensing from *Sodalis praecaptivus* in grain weevils suppresses virulence factors after establishment, allowing for persistent infection (29). Additionally, *Vibrio cholerae* uses quorum sensing to mediate transmission by decreasing biofilm formation to increase dissemination (20, 23). These examples show the complexity of quorum sensing signals inside hosts and highlight the importance of understanding interactions in a host-specific manner.

Autoinducer-2 (AI-2) is a quorum sensing communication molecule that has received a lot of attention because of its effects on both gram-positive and negative bacteria. This molecule is predicted to be produced by 50% of all sequenced bacteria and detected by many more, controlling a broad array of behaviors including virulence, motility, nutrient acquisition, and biofilm formation (22). Because many bacteria produce, sense, and respond to AI-2 signals, quorum sensing via this molecule can result in community level effects. One example is the assembly of multispecies biofilms. In human oral cavities, *Streptococcus gordonii* and *Porphyromonas gingivalis* grow together to form a symbiotic, multispecies biofilm through the production and sensing of this interspecies signaling molecule. In the absence of AI-2, no biofilm is produced; however, biofilm production is restored if either species is able to produce AI-2 (24–26). Also, AI-2 production by gut microbiota may help to mitigate microbial community changes during antibiotic treatment. Mice with a microbiota that produced increased levels of AI-2 maintained a greater diversity of their bacterial gut microbiome after exposure to antibiotics (27). Taken together, these and other studies show that quorum sensing can cause changes in microbial gene expression, bacterial behavior, or community structure, which have resulting effects on host health.

In this study, we present the first analysis of quorum sensing by honey bee associated bacteria. We identified genes encoding the AI-2 producing enzyme (LuxS) in the genomes of both honey bee and bumble bee associated microbes and we predict that this gene is functional by identifying the important catalytic residues. We also provide evidence for the production of the AI-2 molecule within the honey bee gut using a *Vibrio harveyi* biosensor and hypothesize that the hindgut of the honey bee may be a focal chamber for interspecies quorum sensing. We compared newly emerged bees with mature adult worker bees to show that the level of *luxS* expression by one honey bee gut community member (*Gilliamella*) increases as the bacterial community matures. We then used cultured representatives from the major groups associated with the honey bee and identified two clades (*Gilliamella* (Gamma-1), and *Bifidobacteria*) that produce AI-2 when cultured *in vitro*, as expected based on the genomic analysis of LuxS in these genera. Finally, we show that AI-2 can modify an important density-dependent behavior for honey bee host colonization: the formation of a biofilm.

## Methods

### Bioinformatics analysis of luxS in existing genomic datasets

The *luxS* loci were identified based on functional annotations in previously published metagenomic scaffolds (Engel et al. 2012; JGI IMG/M project ID 2498). Scaffolds containing these loci were downloaded from the JGI IMG/M and manually annotated and curated using the Artemis genome browser software (30). Percent identities (amino acid) between LuxS homologs found in the scaffolds were elucidated using National Center for Biotechnology Information’s (NCBI) nucleotide Basic Local Alignment Search Tool (BLAST) suite. We found additional homologs of LuxS in honey bee specific *Gilliamella* and *Bifidobacterium* species through sequence homology with the type strains (accessions NZ_CP007445.1 and NC_018720.1 respectively) using NCBI’s BLAST. All amino acid sequences were downloaded from the NCBI and aligned using MUSCLE with default parameters (31). The alignment was converted to relaxed phylip format and RAxML was used to generate the phylogeny (raxmlHPC-SSE3-m PROTCATBLOSUM62) (32).

### Isolation and culture of bacteria from honeybee guts

Forager honey bees, identified by the presence of provisions in their pollen baskets, were collected via aspirator from colonies maintained at Indiana University – Bloomington. Entire digestive tracts were removed by dissection (the crop, midgut, and hindgut), whole guts were homogenized in sterile PBS, and a dilution series was plated on Brain-Heart Infusion (BHI) agar. Plates were grown anaerobically (via GasPak) at 37°C for 2 days. Isolated colonies were subcultured on BHI to obtain and maintain a pure culture. After 2 days of growth, DNA was extracted using the DNeasy Blood and Tissue kit (QIAGEN), the 16S rRNA was amplified with 27F/1492R primers and sequenced using the 27F primer. Sequences were trimmed for quality and cover > 400bp. Percent identities to known representative strains of honey bee gut microbes determined using NCBI’s BLAST.

### Phylogenetic analysis

From these isolated and taxonomically characterized bacterial cultures, we chose a representative sample that phylogenetically clade with known honey bee specific phylotypes (following (12)). An alignment was generated using 16S rRNA gene sequences and the SINA aligner, which takes into account the 16S rRNA structure. This alignment was used as input to R (version 3.3.3) to create a phylogenetic tree using maximum likelihood with the *ape* and *phangorn* packages. Bootstrap values were generated from 1000 replications.

### Detection of AI-2 via Vibrio harvei reporter

The assay to detect AI-2 production was performed as previously described using a *Vibrio harveyi* reporter strain (34). The *Vibrio harveyi* TL26 reporter strain (*ΔluxN ΔluxS ΔcqsS*; (35)) was used in combination with a positive control: *V. harveyi* BB120 (Wild type). *Vibrio* cultures were grown in autoinducer bioassay (AB) medium aerobically at 30°C overnight. Entire digestive tracts were removed from foragers by dissection and were homogenized in sterile PBS either by section or the entire tract. Honey bee bacterial isolates were grown anaerobically in BHI broth at 37°C for two days. For use in the assay, an overnight culture of TL26 strain was diluted to 1:1000 and 1:5000. Isolates and gut homogenates were tested in triplicate by adding the undiluted, cell free supernatant of each isolate to the 1:5000 dilution of TL26. The positive control contained the cell-free supernatant of BB120 and the 1:5000 dilution of TL26. Cell free supernatant were obtained by centrifuging one mL of culture for five minutes at 14K g. Negative controls contained sterile BHI media and the 1:1000 dilution of TL26. Cell controls contained sterile H_2_O and the 1:1000 dilution of TL26. The plate was incubated at 30°C shaking aerobically for 8 hours. After incubation, luminescence and OD600 were measured on a Synergy H1 plate reader (BioTek Systems). Final luminescence values of the isolates are normalized to the final OD600 of the isolate.

### Detection of Gilliamella apicola LuxS Expression using quantitative RT-PCR analysis

Whole gut sections of newly emerged honey bees as well as adult bees (aged 3-12 days) were collected from established, healthy hives located at Wellesley College in Wellesley, Massachusetts. RNA was extracted using TRIzol reagent and quantitative RT-PCR analyses were performed using SensiFAST™ SYBER Hi-ROX One-Step (Bioline) with primers specific to Gamma-1 LuxS gene (Forward: TTGTATGCCAACACTGTCCTTT, Reverse: TGGCGCGATGATCTTAATTT) and the host actin gene (Forward: ATAGCCAAAACCATGGCAAC, Reverse: TAAAAACCAGTTCGGCAACC, (36)) using the Applied Bioscience StepOnePlus qRT-PCR machine (Life Technologies). Specificity was determined by Sanger sequencing of the amplified product and using NCBI’s BLAST to identify. The expression levels were normalized to the host actin gene using the ΔCt Method.

### Amplicon sequencing of the bacterial community in newly emerged and adult bees

Using the same samples above, RNA was extracted from the homogenates using TRIzol reagent (Ambion). RNA was DNase treated (DNA I, New England Biolabs) and cDNA was synthesized using the SuperScript III First-Strand synthesis system (Invitrogen). cDNA from each sample was amplified via PCR using Earth Microbiome barcoded primers 515F and 806R (37). Earth Microbiome amplification protocols were followed, except for the polymerase used (HF Phusion, New England Biolabs) and amplicons were cleaned with a PCR cleanup kit (Qiagen). Picogreen protocol was used to quantify DNA concentration for each pool sample. Samples were then normalized and pooled collectively for sequencing. Sequencing was performed on an Illumina Miseq, using 250 PE cycles. Sequences are available to reviewers upon request and are currently being deposited at the DDBJ.

### Sequence Analysis

All sequence processing was performed using the Mothur microbial ecology suite (38). Reads from each sample were combined into contiguous sequences and screened for quality (maxambig 0, maxlength 300). Sequences were then aligned with the Silva reference database (silva.bacteria.fasta), preclustered, and examined for chimeras via the uchime function. After removal of chimeric sequences, sequences were taxonomically classified using a honey bee specific training set as a reference (39) and binned into operational taxonomic units (OTUs) based upon 97% sequence identity. The data set was also subsampled to the smallest sample size of 1230 sequences, in order to normalize across samples.

### Biofilm production response to AI-2

To determine if autoinducers (AI) have an effect on biofilm formation we modified a common biofilm assay (40). Using the same isolates used for the AI-2 assays, cultures in early exponential phase were added to a 24-well culture plate (Falcon). For the exogenous autoinducer treatment, a final concentration of 10 µM of AI-2 was added (OMM Scientific). Sterile media was used for the negative control. After a 24-hour aerobic incubation at 37**°**C, crystal violet was added to stain the biofilm. The biofilm was then disrupted with acetic acid and the amount of stain was quantified using absorbance at 600 nm on a Synergy H1 plate reader (BioTek Systems).

## Results

### LuxS gene found in honey bee specific bacterial genomes

LuxS is required for the enzymatic synthesis of the AI-2 signaling molecule. This metalloenzyme, S-Ribosylhomocysteinase, cleaves thioether bonds in S-ribosylhomocysteine resulting in a homocysteine and 4,5-dihydroxy-2,3-pentanedione (DPD) which is the precursor for AI-2. DPD can then spontaneously cyclize to form a furone known as AI-2. These reactions are catalyzed by a divalent metal ion, Fe^2+^, and the activity requires three conserved residues (His-54, His-58, and Cys-126) (21, 41–43).

Using functional annotation of metagenomics scaffolds, the *luxS* gene was identified in the genomes of two honey bee specific microbes (*Gilliamella apicola* and *Bifidobacterium sp.*) (Figure 1A). To determine how conserved this locus was within these bacterial species, we performed a search for homologs, using NCBI’s BLAST, and were able to identify a *luxS* homolog in 91 *Bifidobacterium* species, including both honey bee and bumble bee associated strains (*Bifidobacterium asteroides* and *bombi)*. Similarly, we found *luxS* within 41 *Orbus-*related species, colonizing both honey bees and bumble bees (including *Frischella perrara, Gilliamella apicola*, and *Schmidhempelia bombi*, Figure 1. Finally, alignment of the LuxS protein sequences identified the conserved domains and residues known to be important for LuxS activity (Figure 1C). Highly conserved regions included the catalytic active residue (the cysteine at position 87, arrowhead in Figure 1C) and known metal co-factor binding sites (Asterisks in Figure 1C). This analysis suggested that these LuxS proteins might be both conserved within two important honey bee gut symbiont groups (the *Bifidobacteria* and *gamma-proteobacteria*), and potentially functional.

**Figure 1.**
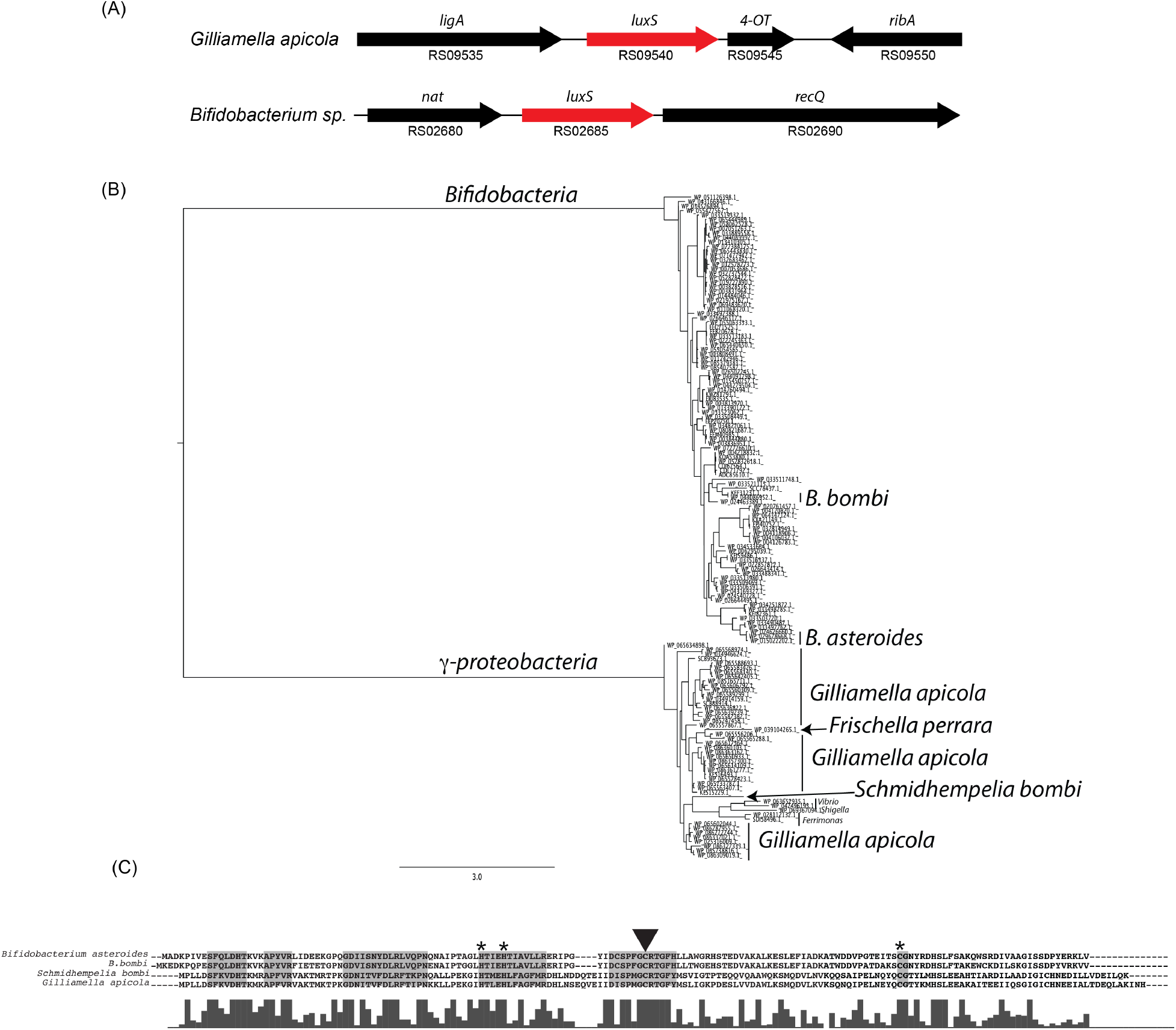
Honey bee associated microbes encode LuxS. **(A)** The *luxS* gene and its syntenic region is shown within the genomes of two honey bee specific isolates (*Gilliamella apicola* and *Bifidobacterium sp.).* **(B)** A phylogeny generated based on aligned LuxS amino acid sequences from honey bee and bumble bee associated microbes. **(C)** LuxS homologs from honey bee specific isolates (*Gilliamella apicola and Bifidobacterium asteroids)* and bumble bee isolates (*Schmidhempelia bombi and Bifidobacterium bombi)*, were identified by functional gene annotation in an existing metagenomic dataset. Shaded areas represent highly conserved regions among these sequences as well as those of other published LuxS homologs (36, 38, 48). Asterisk = conserved iron binding sites; arrowhead = catalytic cysteine.

### Detection of AI-2 production honey bee microbiota

One approach to determine whether LuxS is functional in the honey bee gut symbionts is to identify the production of autoinducer-2 (AI-2). Towards that end, we used a biological reporter assay, where a strain of *Vibrio harveyi* (TL26), which is incapable of producing autoinducers and responds to exogenous AI-2 only, was cultured in the presence of honey bee gut extracts. When TL26 senses AI-2, it responds with the production of light, which we detected using a spectrophotometer (see methods for more detail). We were able to detect significant AI-2 production in entire digestive tracts of honey bee workers as well as gut sections (fore, mid, and hindgut) (Figure 2). These data suggest that AI-2 is being produced within honey bee gut digestive tract.

**Figure 2.**
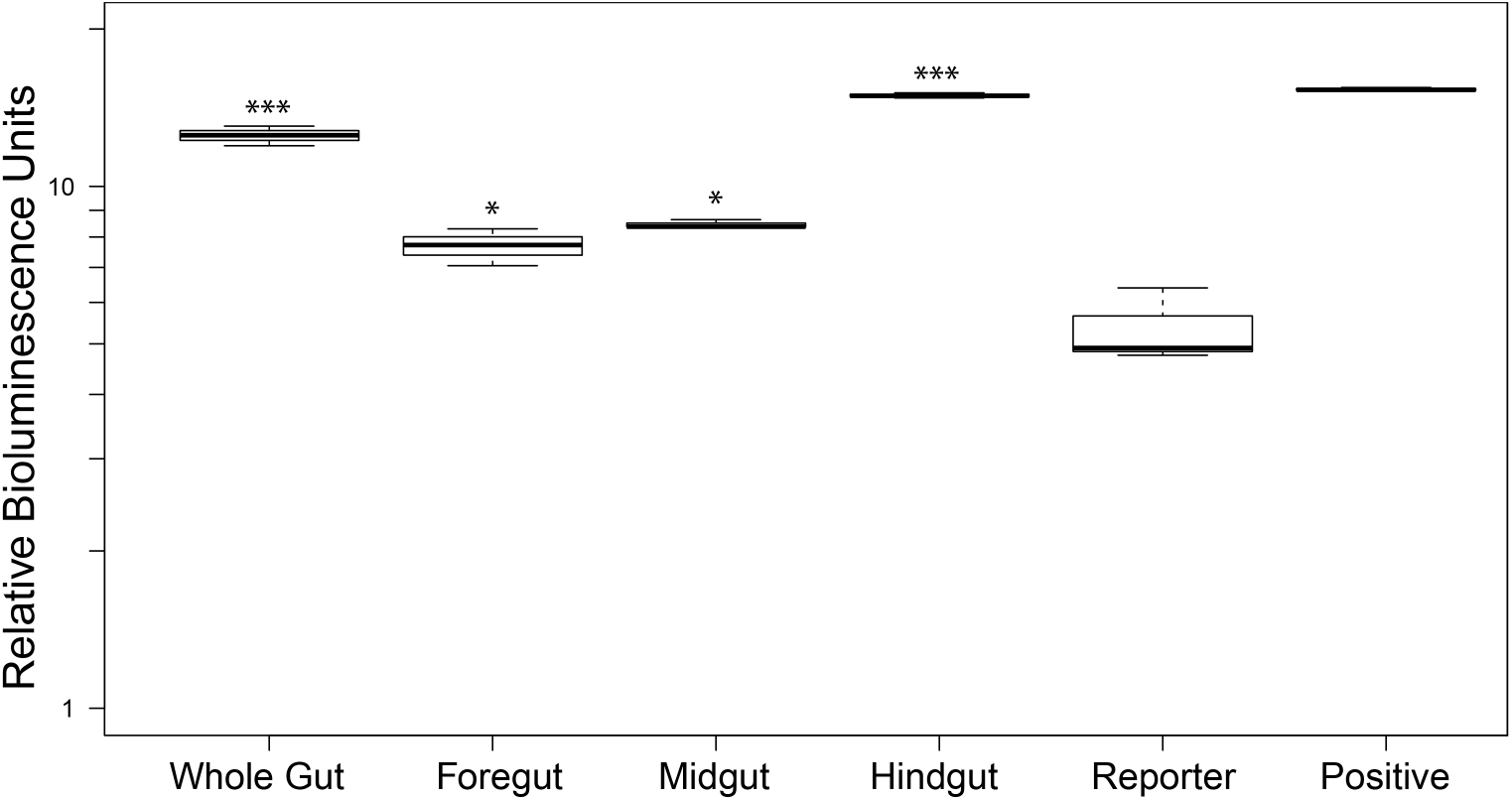
Detection of autoinducer-2 in the honey bee gut. After dissection, entire digestive tracts (Whole Gut) or gut sections (Foregut, Midgut, Hindgut) were homogenized and extracts were used in an autoinducer bioluminescence assay. The production of luminescence by *V. harveyi* TL26 is only observed in the presence of supernatants from the positive control (AI-2 producing *V. harveyi* strain BB120) or from extracts from the honey bee. Note log scale. Samples were compared to reporter alone with a t-test and significance designated by *** = 0.001, ** = 0.01, * = 0.05.

### Gilliamella strains express luxS in mature adult bees

Because our assay above included honey bee tissue, we sought to confirm that the AI-2 signal we observed was coming from the bacterial community members. We therefore marked, age matched, and collected newly emerged bees (< 1 day old) and mature adult worker bees (> 3 days old). After extracting RNA from these samples, we used qRT-PCR and a *Gilliamella-*specific *luxS* primer set to quantify the expression of the *luxS* gene relative to host actin. In addition, we also characterized the microbial community associated with these same samples to confirm the establishment of *Gilliamella* in the mature adult worker bees. Previous work suggests that newly emerged bees lack the characteristic gut microbiome found in adult worker bees (18) and we confirmed that our newly emerged bee samples also lacked the core community and instead were dominated by unclassified OTUs (Figure 3). In contrast, mature adult bees were colonized by the characteristic core community (Figure 3). Additionally, *luxS* expression by *Gilliamella* was not detected in newly emerged bees while we identified low, but consistent expression in mature adult bees (relative expression compared to host actin was 4.40E-05 +/-1.35E-05 SE; Figure 3).

**Figure 3.**
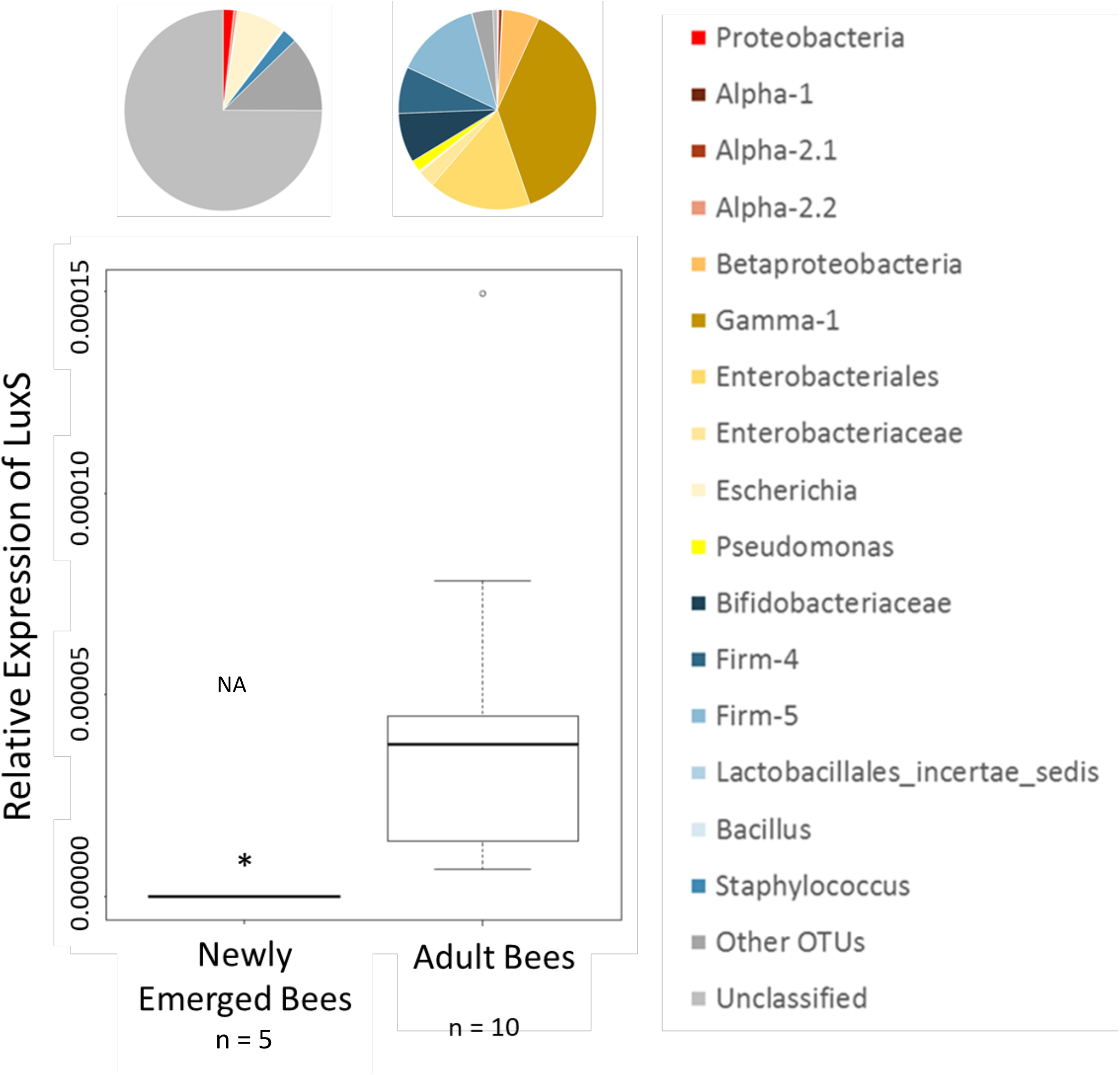
*luxS* gene expression increases and bacterial community composition changes as adult bees mature. Bacterial community composition, based on 16S rRNA, in newly emerged bees is dominated by unclassified bacterial taxa whereas adult bees have acquired the characteristic worker bee microbiome. Additionally, relative expression of *luxS* (qPCR) by *Gilliamella apicola* is detectable in mature adult bees while in newly emerged bees we observed no amplification of the transcript (NA = No Amplification).

### Gilliamella *and* Bifidobacteria *species* produce AI-2 in vitro

Our results using gut extracts suggested that the honey bee gut community members may be producing AI-2 *in vivo.* To support our qRT-PCR results and the bioinformatics analysis of the *luxS* locus, we cultured *Gilliamella* and *Bifidobacteria* species from the honey bee gut and subjected their supernatants to the *Vibrio harveyi* AI-2 reporter assay. Representative isolates from the prominent clades found in the honey bee were chosen for the assay based on their phylogenetic placement (based on 16S rRNA gene sequence; Figure 4A). Each chosen isolate formed a highly supported clade (100% confidence) with known, characterized honey bee microbiome members (Figure 4A). In addition, 16S rRNA sequences from the cultured isolates were 92% - 100% identical to known honey bee associated microbes (Figure 4B). After normalizing to the optical density of the cultures, we were able to detect AI-2 production by strains from each of these genera (as inferred from the luminescence produced by TL26) (Figure 4C). Compared to the negative controls, we observed statistically significant luminescence by TL26 in the presence of supernatants from *Bifidobacteria* and *Gilliamella* species (Figure 4C).

**Figure 4.**
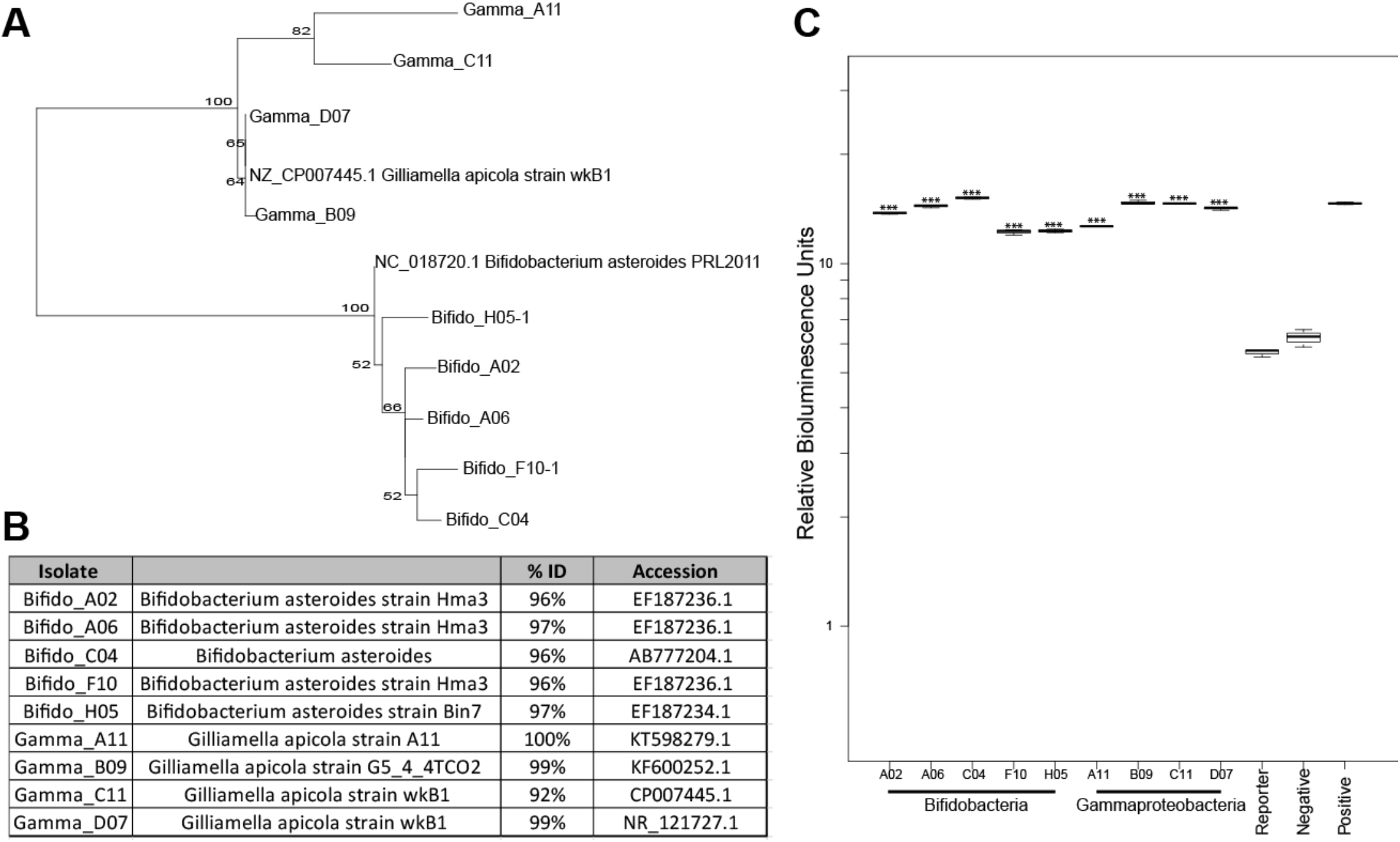
*Gilliamella apicola* and *Bifidobacterium sp.* produce AI-2 *in vitro*. **(A)** Phylogenetic tree of bacterial isolates utilized in this study and their evolutionary placement in the context of other honey bee gut microbes. 16S rRNA genes (> 400 bp) were used to construct this maximum likelihood phylogeny and bootstrap values are from 1000 iterations **(B)** 16S rRNA gene sequences from cultured isolates are 92-100% identical to known honey bee associated microbes. Percent identities of cultured isolates shown relative to accessions in the NCBI’s nr database. **(C)** Detection of AI-2 in the overnight culture supernatants of *Gilliamella apicola* and *Bifidobacterium sp.* using *Vibrio harveyi* reporter strain luminescence. Controls (negative: sterile BHI; positive: *V. harveyi* BB120) (Note log scale). Samples compared to the Reporter only control with a t-test and significance designated by *** = < 0.001.

### Biofilm production is modulated by honey bee bacterial isolates in response to autoinducers

Previous work had identified the presence of a microbial biofilm in the honey bee digestive tract. Because biofilm production is known to be regulated by quorum sensing, we sought to identify a relevant and functional link between the production of AI-2 in the honey bee and host colonization. With the same representative isolates used above, we cultured these microbes with or without exogenously added AI-2. We identified a statistically significant increase in biofilm production with added AI-2 for all our four isolates from the *Gilliamella* genus (Figure 5). However, there were no significant changes in the biofilm production of the *Bifidobacteria* isolates in response to the addition of AI-2 (Figure 5).

**Figure 5.**
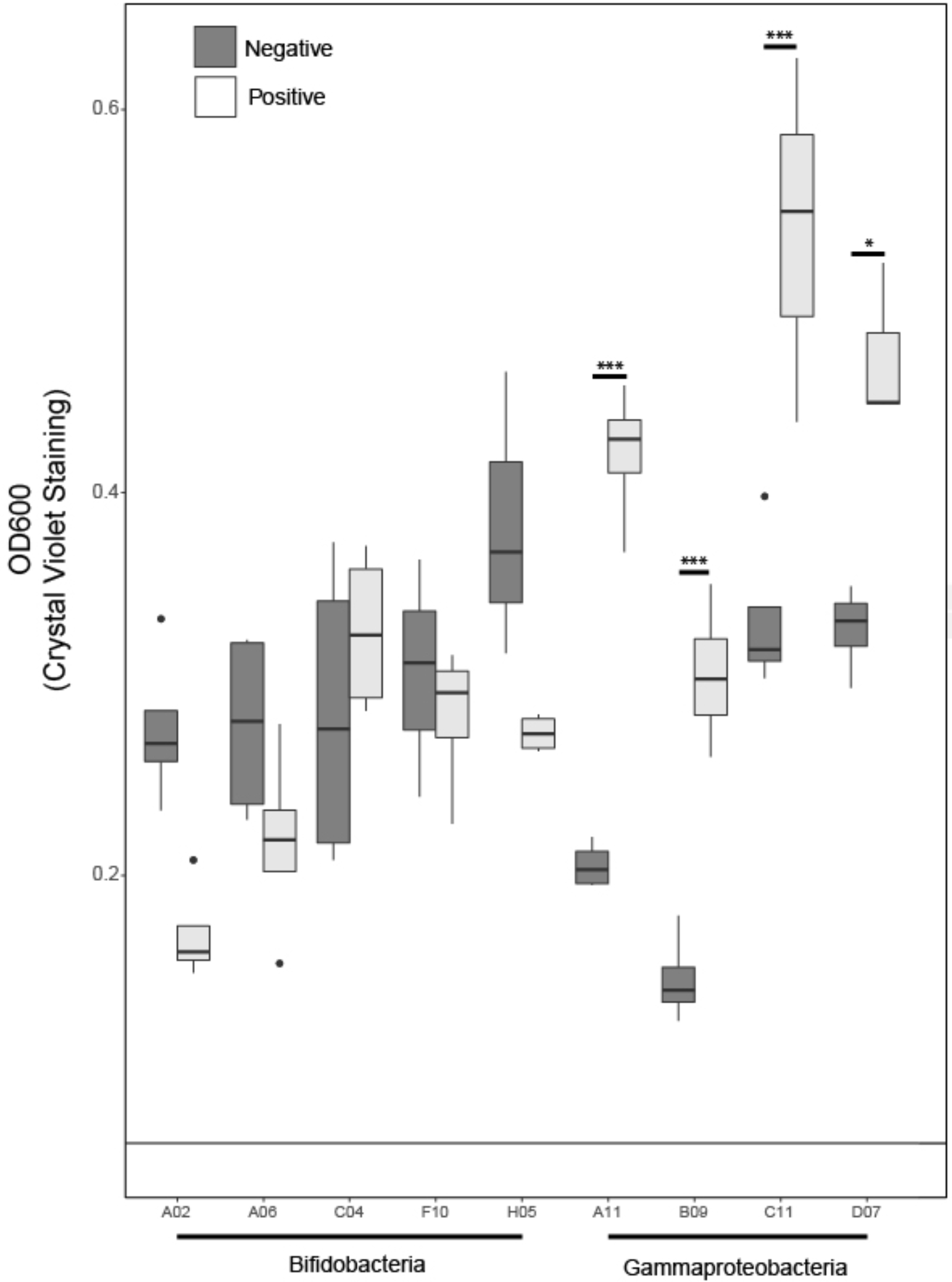
Quantification of crystal violet stained biofilms produced by honey bee gut microbiome members. Biofilm production on a chitin substrate by honey bee associated microbes was quantified using a standard crystal violet assay. Cultures were incubated either without (dark grey) or with (light grey) purified AI-2 added (see methods). Black line across the graph represents the average absorbance from sterile media controls across treatments. Comparisons were made between isolates treated with AI-2 and the same isolate without AI-2 added using t-tests. Significance designated by *** = < 0.001, * = p < 0.05.

## Discussion

Bacterial infections of eukaryotic hosts are established and maintained using a variety of bacterial behaviors such as biofilm formation, motility, and virulence. These behaviors are often modulated and controlled in a density dependent fashion using signaling molecules (44). Here we present multiple lines of evidence to support bacterial autoinducer-2 based quorum sensing in the honey bee microbiota. We identified an open reading frame, annotated as encoding LuxS, the AI-2 producing enzyme, in honey bee specific isolates. We detected the production of AI-2 *in vivo* using whole bee gut extracts. Additionally, we showed that *luxS* expression by *Gilliamella* is only detected in adult bees that harbor the core bacterial community. Finally, we also demonstrated that specific isolates, representative of the honey bee core microbiome, produce AI-2 and increase biofilm formation in response to AI-2. These data support our conclusion that honey bee associated bacteria produce AI-2 during colonization of the host.

The two genera we worked with here (*Bifidobacteria* and *Gilliamella*) have been implicated in honey bee health or nutrition. For example, lactic acid bacteria (*Bifidobacteria*) isolated from the honey bee crop have been shown to protect larvae from pathogens such as European Foulbrood, likely through the production of antimicrobial molecules (45, 46). Members from the *Gilliamella* likely contribute to degradation of plant carbohydrates as they degrade pectin *in vitro* (15). Therefore, our work identifies a potential mechanism by which functionally important members of the honey bee microbiota may communicate with each other during their host association and suggests that AI-2 may regulate density dependent behaviors, such as biofilm formation, in the honey bee microbiota. In fact, *Gilliamella* species are known to form a multispecies biofilms in the honey bee digestive tract (18). For example, the ileum is colonized by both Gamma- and Beta-proteobacteria and the rectum is dominated by Firmicutes and Gamma-proteobacteria (18). These stratified biofilms suggest that colonization dynamics or environmental gradients may play a role in the colonization of the host. While important for the establishment and persistence of the microbiota, biofilms may also facilitate the breakdown of plant materials in host digestion. Based on our data, we propose that the production of autoinducers may mediate the colonization of honey bee specific microbes, contributing to the stratified biofilm observed. While some *Bifidobacterium* isolates were able to form a biofilm, the production of which was unaffected by exogenous AI-2, all *Gilliamella* strains increased their biofilm production in the presence of added AI-2. We therefore hypothesize that species such as *Bifidobacterium* may colonize early and that production of AI-2 by these early colonizers may allow other species to form biofilms at higher density. This hypothesis awaits further testing.

The presence of *luxS* in *Frischella* as well as isolates from *Bombus* species suggests that social behaviors in these microbes, such as the production of AI-2, may be conserved across bee associated microbes, both pathogens and mutualists. The production of AI-2 and quorum sensing *writ large* is not uniformly beneficial to a host, as virulence is another density dependent behavior often regulated by quorum sensing. For example, in the cabbage white butterfly (CWB), quorum sensing in pathogenic *Pseudomonas aeruginosa* contributes to virulence such that, when quorum sensing pathways are disrupted, CWB larval survival rates are increased (49). Similarly in mammals, *Pseudomonas aeruginosa* utilizes quorum sensing during chronic lung infections, upregulating biofilm formation and adhesion (50, 51). Importantly, we identified a *luxS* in *Frischella perrara*, a putative bee pathogen, and in this organism, AI-2 may be utilized to promote pathogenicity within the honey bee host (52).

To our knowledge, this is the first time quorum sensing has been shown to occur in the honey bee microbiota. Investigations such as this one can help to identify not only behaviors mediated by quorum sensing but potential cross-talk and communication *between* microbial members in the gut. For example, although we focused on a single quorum sensing molecule (AI-2), there are likely many other molecules (such as AHLs and oligopeptides) produced *in vivo* by honey bee gut microbes. To form a complete picture of microbial communication between community members, additional quorum sensing molecules need be examined as well as their effects on gene regulation. We know that the honey bee bacterial community is specific and consistent (in terms of the presence of members), however the proportion of different bacteria within individual bees can vary (16, 19). If these bacterial members are participating in intra-species communication and mediating important behaviors, their relative proportions may impact other community members and potentially host health. Future work is needed to understand how these persistent infections are maintained and influenced by quorum sensing.

## Acknowledgements

We thank Dr. Julia van Kessel for generous gifts of *Vibrio harveyi* biosensor strains and synthesized AI-2. We thank Newton lab members Delaney Miller and Eric Smith for feedback in initial stages of writing this manuscript. This project was funded, in part, by crowd sourcing through experiment.com. ILGN was supported by generous startup funds from Indiana University and KIM was supported by the Floyd Microbiology Fellowship.

## References

1. Lee W-J, Brey PT. 2013. How Microbiomes Influence Metazoan Development:Insights from History and Drosophila Modeling of Gut-Microbe Interactions. Annu Rev Cell Dev Biol 29:571–592.

2. Shin SC, Kim S-H, You H, Kim B, Kim AC, Lee K-A, Yoon J-H, Ryu J-H, Lee W-J. 2011. Drosophila Microbiome Modulates Host Developmental and Metabolic Homeostasis via Insulin Signaling. Science (80-) 334:670–4.

3. Sharon G, Segal D, Ringo JM, Hefetz A, Zilber-Rosenberg I, Rosenberg E. 2010. Commensal bacteria play a role in mating preference of Drosophilia melanogaster. Proc Natl Acad Sci 107:20051–20056.

4. Weiss B, Aksoy S. 2011. Microbiome influences on insect host vector competence. Trends Parasitol 27:514–522.

5. Raymann K, Shaffer Z, Moran NA. 2017. Antibiotic exposure perturbs the gut microbiota and elevates mortality in honeybees. PLoS Biol 15:1–22.

6. Jones RT, Knight R, Martin AP. 2010. Bacterial communities of disease vectors sampled across time, space, and species. ISME J 4:223–231.

7. Rosenthal GA, Berenbaum MR. 1992. Herbivores: Their Interactions with Secondary Plant Metabolites, Second Edition: Ecological and Evolutionary Processes. Academic Press.

8. Klein A-M, Vaissiere BE, Cane JH, Steffan-Dewenter I, Cunningham SA, Kremen C, Tscharntke T. 2007. Importance of pollinators in changing landscapes for world crops. Proc R Soc B Biol Sci 274:303–313.

9. Jonsson M, Malmqvist B. 2003. Mechanisms behind positive diversity effects on ecosystem functioning: testing the facilitation and interference hypotheses. Oecologia 134:554–559.

10. Jones CG, Lawton JH, Shachak M. 1994. Organisms as Ecosystem Engineers Organisms as ecosystem engineers. Oikos 69:373–386.

11. Mattila HR, Rios D, Walker-Sperling VE, Roeselers G, Newton ILG, V.E. W-S, Roeselers G, Newton ILG. 2012. Characterization of the Active Microbiotas Associated with Honey Bees Reveals Healthier and Broader Communities when Colonies are Genetically Diverse. PLoS One 7:e32962.

12. Newton ILG, Roeselers G. 2012. The effect of training set on the classification of honey bee gut microbiota using the Naive Bayesian Classifier. BMC Microbiol 12:221.

13. Moran NA, Hansen AK, Powell E, Sabree ZL. 2012. Distinctive gut microbiota of honey bees assessed using deep sampling from individual worker bees. PLoS One 7:e36393.

14. Martinson VG, Danforth BN, Minckley RL, Rueppell O, Tingek S, Moran NA. 2011. A simple and distinctive microbiota associated with honey bees and bumble bees. Mol Ecol 20:619–628.

15. Engel P, Martinson VG, Moran NA. 2012. Functional diversity within the simple gut microbiota of the honey bee. PNAS www.pnas.o.

16. Lee FJ, Rusch DB, Stewart FJ, Mattila HR, Newton ILG. 2015. Saccharide breakdown and fermentation by the honey bee gut microbiome. Environ Microbiol 17:796–815.

17. Kesnerova L, Mars RAT, Ellegaard KM, Troilo M, Sauer U, Engel P. 2017. Disentangling metabolic functions of bacteria in the honey bee gut. doi.org.

18. Martinson VG, Moy J, Moran N a. 2012. Establishment of characteristic gut bacteria during development of the honeybee worker. Appl Environ Microbiol 78:2830–2840.

19. Rokop ZP, Horton MA, Newton ILG. 2015. Interactions between Cooccurring Lactic Acid Bacteria in Honey Bee Hives. Appl Environ Microbiol 81:7261–7270.

20. Waters CM, Bassler BL. 2005. Quorum Sensing: Cell-to-Cell Communication in Bacteria. Annu Rev Cell Dev Biol 21:319–346.

21. Miller MB, Bassler BL. 2001. Quorum Sensing in bacteria. Annu Rev Microbiol 55:165–199.

22. Federle MJ, Bassler BL. 2003. Interspecies communication in bacteria. J Clin Invest 112:1291–1299.

23. Hammer BK, Bassler BL. 2003. Quorum sensing controls biofilm formation in Vibrio cholerae. Mol Microbiol 50:101–114.

24. Pereira CS, Thompson JA, Xavier KB. 2013. AI-2-mediated signalling in bacteria. FEMS Microbiol Rev 37:156–181.

25. McNab R, Ford SK, El-sabaeny A, Barbieri B, Cook GS, Lamont RJ. 2003. LuxS-Based Signaling in Sfilm Formation with Porphyromonas gingivalis. J Bacteriol 185:274–284.

26. Rickard AH, Palmer RJ, Blehert DS, Campagna SR, Semmelhack MF, Egland PG, Bassler BL, Kolenbrander PE. 2006. Autoinducer 2: A concentration-dependent signal for mutualistic bacterial biofilm growth. Mol Microbiol 60:1446–1456.

27. Thompson JAA, Oliveira RAA, Djukovic A, Ubeda C, Xavier KBB. 2015. Manipulation of the Quorum Sensing Signal AI-2 Affects the Antibiotic-Treated Gut Microbiota. Cell Rep 10:1861–1871.

28. Nyholm S V., McFall-Ngai M. 2004. The winnowing: establishing the squid–vibrio symbiosis. Nat Rev Microbiol 2:632–642.

29. Enomoto S, Chari A, Clayton AL, Dale C. 2017. Quorum Sensing Attenuates Virulence in Sodalis praecaptivus. Cell Host Microbe 21:629–636.e5.

30. Rutherford K, Parkhill J, Crook J, Horsnell T, Rice P, Rajandream M a, Barrell B. 2000. Artemis: sequence visualization and annotation. Bioinformatics 16:944–945.

31. Edgar RC. 2004. MUSCLE: Multiple sequence alignment with high accuracy and high throughput. Nucleic Acids Res 32:1792–1797.

32. Stamatakis A. 2014. RAxML version 8: A tool for phylogenetic analysis and post-analysis of large phylogenies. Bioinformatics 30:1312–1313.

33. Tamura K, Stecher G, Peterson D, Filipski A, Kumar S. 2013. MEGA6: Molecular Evolutionary Genetics Analysis Version 6.0. Mol Biol Evol 30:2725–2729.

34. Surette MG, Bassler BL. 1998. Quorum sensing in Escherichia coli and Salmonella typhimurium. Proc Natl Acad Sci U S A 95:7046–7050.

35. Long T, Tu KC, Wang Y, Mehta P, Ong NP, Bassler BL, Wingreen NS. 2009. Quantifying the Integration of Quorum-Sensing Signals with Single-Cell Resolution. PLoS Biol 7:e68.

36. vanEngelsdorp D, Evans JD, Saegerman C, Mullin C, Haubruge E, Nguyen BK, Frazier M, Frazier J, Cox-Foster D, Chen Y, Underwood R, Tarpy DR, Pettis JS. 2009. Colony collapse disorder: A descriptive study. PLoS One 4.

37. Caporaso JG, Lauber CL, Walters WA, Berg-Lyons D, Huntley J, Fierer N, Owens SM, Betley J, Fraser L, Bauer M, Gormley N, Gilbert JA, Smith G, Knight R. 2012. Ultra-high-throughput microbial community analysis on the Illumina HiSeq and MiSeq platforms. Isme J 6:1621–1624.

38. Kozich JJ, Westcott SL, Baxter NT, Highlander SK, Schloss PD. 2013. Development of a dual-index sequencing strategy and curation pipeline for analyzing amplicon sequence data on the MiSeq Illumina sequencing platform. Appl Env Microbiol2013/06/25. 79:5112–5120.

39. Newton ILG, Roeselers G. 2012. The effect of training set on the classification of honey bee gut microbiota using the Naive Bayesian Classifier. Bmc Microbiol 12.

40. O’Toole GA. 2011. Microtiter Dish Biofilm Formation Assay. J Vis Exp 3–5.

41. Zhu J, Patel R, Pei D. 2004. Catalytic mechanism of S-ribosylhomocysteinase (LuxS): Stereochemical course and kinetic isotope effect of proton transfer reactions. Biochemistry 43:10166–10172.

42. Pei D, Zhu J. 2004. Mechanism of action of S-ribosylhomocysteinase (LuxS). Curr Opin Chem Biol 8:492–497.

43. Hilgers MT, Ludwig ML. 2001. Crystal structure of the quorum-sensing protein LuxS reveals a catalytic metal site. Proc Natl Acad Sci U S A 98:11169–11174.

44. Williams P, Winzer K, Chan WC, Camara M. 2007. Look who’s talking: communication and quorum sensing in the bacterial world. Philos Trans R Soc B Biol Sci 362:1119–1134.

45. Vásquez A, Forsgren E, Fries I, Paxton RJ, Flaberg E, Szekely L, Olofsson TC. 2012. Symbionts as Major Modulators of Insect Health: Lactic Acid Bacteria and Honeybees. PLoS One 7:e33188.

46. Forsgren E, Olofsson TC, Vásquez A, Fries I. 2009. Novel lactic acid bacteria inhibiting Paenibacillus larvae in honey bee larvae. Apidologie 41:99–108.

47. Martinson VG, Moy J, Moran NA. 2012. Establishment of Characteristic Gut Bacteria during Development of the Honeybee Worker. Appl Environ Microbiol 78:2830–2840.

48. Martinson VG, Danforth BN, Minckley RL, Rueppell O, Tingek S, Moran NA. 2011. A simple and distinctive microbiota associated with honey bees and bumble bees. Mol Ecol 20:619–628.

49. Borlee BR, Geske GD, Robinson CJ, Blackwell HE, Handelsman J. 2008. Quorum-sensing signals in the microbial community of the cabbage white butterfly larval midgut. ISME J 2:1101–11.

50. Davies DG, Parsek MR, Pearson JP, Iglewski BH, Costerton JW, Greenberg EP. 1998. The involvement of cell-to-cell signals in the development of a bacterial biofilm. Science 280:295–298.

51. Smith RS, Iglewski BH. 2003. P. aeruginosa quorum-sensing systems and virulence. Curr Opin Microbiol 6:56–60.

52. Emery O, Schmidt K, Engel P. 2017. Immune system stimulation by the gut symbiont Frischella perrara in the honey bee (Apis mellifera). Mol Ecol 26:2576–2590.

